# DAB2IP inhibits p53 ubiquitin-mediated degradation by competitively binding to GRP75 and suppresses tumor malignancy in colon cancer

**DOI:** 10.1101/2021.06.28.450115

**Authors:** Jie Shen, Shengjie Feng, Jiao Deng, Qingwen Huang, Dayong Zhao, Weiyi Jia, Xiaolan Li, Deding Tao, Jianping Gong, Daxing Xie, Liang Liu

## Abstract

Increasing evidence has shown that DAB2IP acts as a tumor suppressor and plays an inhibition role in many tumors. However, the underlying mechanism is still uncertain. Our study shows that DAB2IP is positively associated with a better prognosis in colon cancer patients with wild-type TP53 expression. In vitro assay shows that DAB2IP elicits potent tumor-suppressive effects on inhibiting cell invasiveness, colony formation and promoting cell apoptosis in wild-type TP53 colon cancer cell lines. Subsequently, DAB2IP is demonstrated to up-regulate the stability of wild-type TP53 by inhibiting its degradation in a ubiquitin-proteasome-dependent manner. Using mass spectrometry profiling, we unveil that DAB2IP and p53 could both interact with the ubiquitin ligase-related protein, GRP75. Mechanistically, DAB2IP could competitively bind with GRP75, thus reducing GRP75-mediated p53 ubiquitination and degradation. Finally, animal experiments also reveal that DAB2IP inhibits the tumor progression in vivo. In conclusion, our study presents a novel function of DAB2IP in GRP75-driven wild-type p53 degradation, which provides a new insight in DAB2IP-induced tumor suppression and provides a novel molecular aspect of the p53 pathway.

## Introduction

Colorectal cancer (CRC) is the fourth leading cause of cancer-related deaths worldwide (1). Most CRCs develop from the sequential inactivation of tumor suppressor genes, including APC, TP53, and SMAD4, as well as the activation of oncogenes, including RAS and BRAF (2). TP53 was first discovered and classified as a cellular SV40 large T antigen-binding protein (3). As a classical tumor suppressor, mutations in p53 are frequently observed in multiple cancer types and are critical factors in tumorigenesis (4). Previous studies have revealed that missense mutations in p53 proteins are common in human cancer, and these mutant proteins acquire oncogenic properties that promote proliferation, cell survival and metastasis (5). In colorectal cancer, p53 mutations have been reported to occur in approximately 40%-50% of patients (2, 6). Although the oncological function of mutant p53 has been widely explored, there is still a lack of understanding of the remaining CRCs with wild-type p53.

Wild-type p53 mediates cell cycle arrest and provides a cell death checkpoint, and wild-type p53 can be activated by multiple cellular stresses (7). Functionally, p53 is stabilized and binds to DNA as a tetramer, and results the transcriptional regulation of genes, such as PUMA, BAX, and CDKN1A, that are involved in DNA repair, cell-cycle arrest, senescence and apoptosis in a sequence-specific manner (7, 8). In addition to genetic mutations, emerging studies have also shown that the protein level or activity of endogenous wild-type p53 could be downregulated in a considerable proportion of cancer patients through epigenetic or posttranslational mechanisms (9). Due to the significant role of p53 inactivation in tumorigenesis and therapeutic responses (10–12), it is critical to explore the molecular mechanism by which wild-type p53 is dysregulated in tumors.

DAB2IP, also called ASK1-interacting protein (AIP1), is often reported to act as a tumor suppressor and frequently silenced by epigenetic modifications in many tumors (13–18). As a protein with GTPase activity, DAB2IP is involved in many signaling pathways associated with tumorigenesis, including the Ras-Raf, PI3K-Akt and ASK1-JNK (19–21). Additionally, DAB2IP also presents a function of scaffold protein in modulating nuclear b-catenin/T-cell factor activity, enhancing PROX1 mediated HIF1a ubiquitination, down-regulating NF-κB signaling, thereby inhibit epithelial-mesenchymal transition, cancer stem cell signatures and suppress tumor progression (22–24). A recent study found that mutant p53 potentiates the oncogenic effects of insulin and TNF-α by inhibiting DAB2IP (25, 26). Whether DAB2IP could, in reverse, regulate wild-type p53 in CRC is still not clear.

In this research, we provided evidences that DAB2IP exerted tumor-suppressive effects in wild-type p53 colon cancer, and involved in TP53 degradation in a ubiquitin-proteasome-dependent manner. Furthermore, we unveiled that DAB2IP and p53 could both interact with the ubiquitin ligase-related protein, GRP75. Mechanistically, DAB2IP could competitively bind with GRP75, thus reducing GRP75-mediated p53 ubiquitin-mediated degradation, and finally inhibit tumor progression in colon cancer. Collectively, our study presents a novel function of DAB2IP in GRP75-driven wild-type p53 degradation and reveals a new signaling axis that is involved in DAB2IP-induced tumor suppression in CRC.

### Materials/Subjects and Methods: Reagents and antibodies

Cell culture medium and other supplements were purchased from KeyGEN Biotech (Nanjing, China). Fetal bovine serum was obtained from MULTICELL (Canada). Basic chemical reagents for molecular biology were purchased from Sigma-Aldrich (St. Louis, MO). Antibodies used in this research were described as follow.

| Antibody | Manufacturer | Cat. # |
| --- | --- | --- |
| DAB2IP | Abcam, Inc. (Cambridge, UK) | Ab87811 |
| p53 | Santa Cruz Biotechnology, Inc. (Santa Cruz, CA) | sc-1226 |
| p21 | Cell Signal Technology, Inc. (Beverly, MA) | 2947 |
| BAX | Cell Signal Technology, Inc. (Beverly, MA) | 5023 |
| GRP75 | Cell Signal Technology, Inc. (Beverly, MA) | 3593 |
| GAPDH | Cell Signal Technology, Inc. (Beverly, MA) | 97166 |
| Myc | Cell Signal Technology, Inc. (Beverly, MA) | 2276 |
| HA | Cell Signal Technology, Inc. (Beverly, MA) | 3724 |
| FLAG | Cell Signal Technology, Inc. (Beverly, MA) | 14793 |

### Tissue microarray and Immunohistochemistry (IHC)

Human colon cancer tissue microarray of 78 cases primary lesion and 2 adjacent normal colon tissue (US Biomax, Cat,# CO802c) was used evaluated the expression of DAB2IP and P53. Slide was dewaxed, rehydrated and heated in sodium citrate buffer (0.01 M, pH 6.0) for antigen retrieval. Subsequently, the endogenous peroxidase was inhibited with 3% hydrogen peroxide with 0.1% sodium azide for 30 min and non-specific staining was blocked through incubation in 5% bovine serum albumin for 2 hours. Next, slide was incubated with 1:200 diluted DAB2IP antibodies or 1:400 diluted P53 antibodies at 4°C overnight and with biotinylated secondary antibody for 2 hours. DAB Horseradish Peroxidase Color Development Kit (Wuhan BosterBio Co.Ltd, Cat. #AR1022) was used for Immuno-staining and counterstain was performed by hematoxylin staining. The results were analyzed under a microscope.

The expression level of DAB2IP and P53 was evaluated by IHC score, which was calculated through multiplying proportion score and intensity score, and categorized into level 1 (IHC score 0–3), level 2 (IHC score 4-6) and level 3 (IHC score more than 6). The proportion score reflected the fraction of positive staining cells (0, none; 1, ≤10%; 2, 10% to ≥25%; 3, >25% to 50%; 4, >50%), and the intensity score revealed the staining intensity (0, no staining; 1, weak; 2, intermediate; 3, strong).

### Cell lines and cell culture

Colon cancer cell lines, Caco-2, HT29, HCT116, KM12, Lovo, SW48, SW480, SW620 and human embryonic kidney cell line HEK293T were purchased from American Type Culture Collection (ATCC). All the cell lines were cultured routinely in Dulbecco’s Modified Eagle Medium supplemented with 10% fetal bovine serum and 1% penicillin/streptomycin at 37 °C and in 5% CO2.

### Small interfering RNAs, Plasmids and vectors constructing

Small interfering RNAs (siRNAs) targeting DAB2IP and GRP75 were designed and synthesized by Guangzhou RiboBio Co. Ltd. The sequences of siRNAs were described as follow.

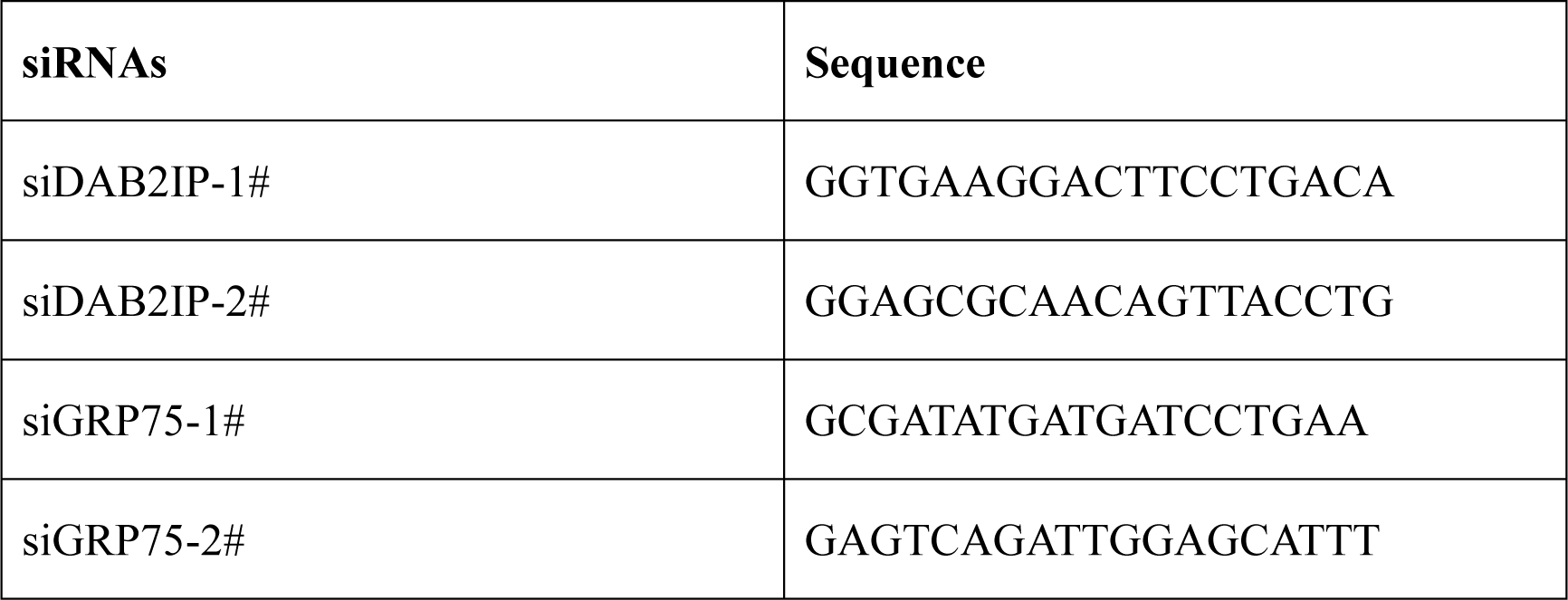

pcDNA3.1-Flag-tagged-DAB2IP, pcDNA3.1-Myc-tagged-DAB2IP overexpressing plasmids were gifts from Jer-Tsong Hsieh. pcDNA3.1-TP53 overexpressing plasmid was generated before and were verified by performing sequencing. GRP75 DNA fragment was obtained by PCR and cloned into pcDNA3.1 vector containing Green fluorescent protein (GFP) tag. HA-ubiquitin plasmid was purchased from Addgene (Cat.#17608). pLKO-DAB2IP-lentiviral-shRNA was constructed previously and used for knocking down DAB2IP expression. The sequences of shDAB2IP were described as follow.

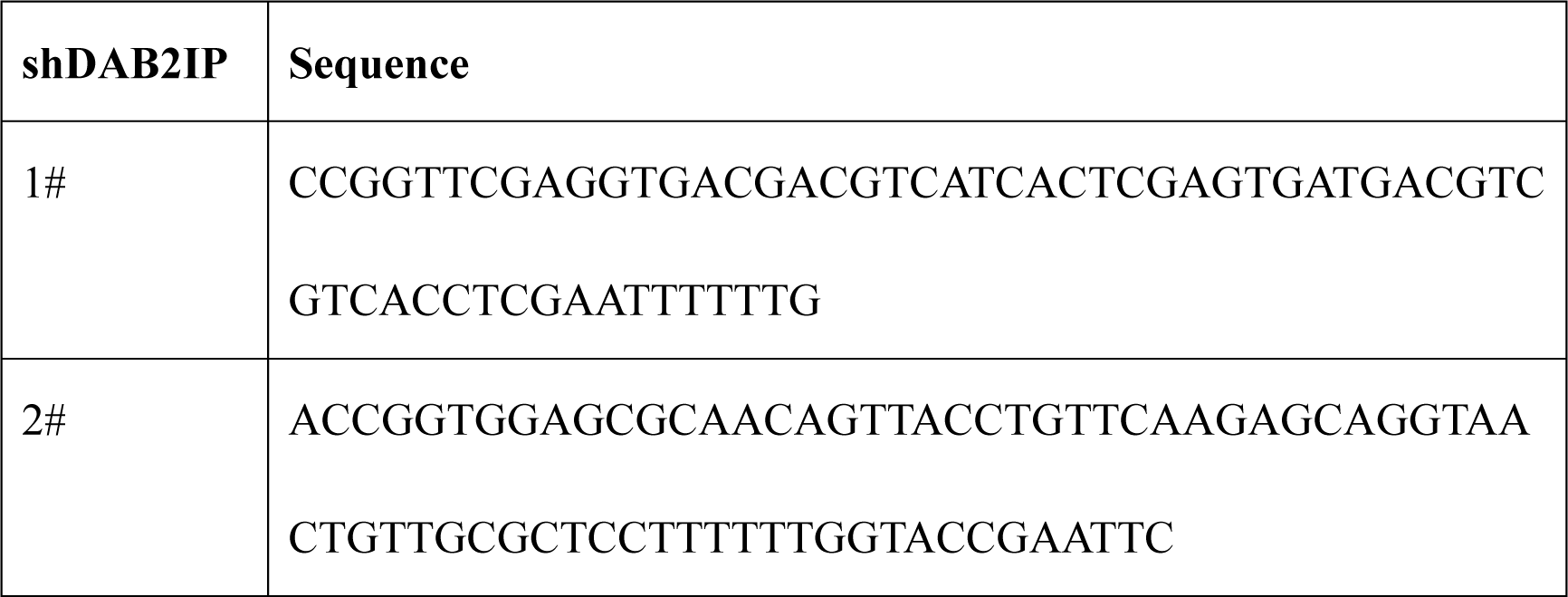

All the plasmids were verified by performing sequencing.

### Transfection and establishment of stable-expressing cell line

Small interfering RNAs transfections were performed using Lipofectamine®2000 transfection reagent (Thermofisher, Cat. #11668019) according to the manufacturès protocol. Non-targeting siRNA (siNC) were used for negative control. Plasmid transfections were performed using Lipofectamine®3000 transfection reagent (Thermofisher, Cat. # L3000015) according to the manufacturès manual. After 48 hours, cell biological and biochemical experiments were performed.

For establishment of stable-expressing cell lines, recombinant lentivirus of sh-scramble, sh-DAB2IP (purchased from Origene) were transfected into HCT116 and SW48 according to the manufacturer’s suggested protocol. After transfection, cells were treated with 10μg/ml puromycin (Thermofisher, Cat. # A1113802) for 4 weeks to get stable cell lines. DAB2IP expression of the stable cell lines was confirmed via western blotting.

### Transwell assay, colony formation assay and apoptosis assay

24-well Transwell plates with pore size 8um (Corning, Cat. #3422) were used for Transwell assay. 1*10^5^ cells were harvested in 100μl of serum free culture-medium and added into the upper chamber. Then 600μl 30% fetal bovine serum medium was placed into the bottom compartment of the chamber as a source of chemo-attractant. After 24 hours culture, cells that crossed the inserts were fixed and strained with microscope (100X magnification).

For the colony-formation assay, the cells were plated in six-well plates at a density of 500 cells/well and were cultured for 2 weeks. Then culture medium was removed. The colonies were fixed in methanol, stained with Crystal-Violet and photographed.

For apoptosis analysis, 1 × 10^4^ cells were seeded on six-well plates and cultured to reach 70% confluence. Then cells were treated with 10ug/ml 5-fluorouracil (5-FU) for 24 hours. Next, cells were collected by 0.02% trypsin without ethylene diamine tetra acetic acid (EDTA) and stained with annexin V-FITC/ propidium iodide kit (BD Cat. #559763) according to the manufacturer’s manual, and analyzed by flow cytometer.

### Western blot assay and immunoprecipitation

Total protein was extracted via NP40 lysis buffer added with Phenylmethylsulfonyl fluoride (PMSF) and cocktail protease inhibitor (MedChemExpress, Cat. # HY-K0010) and separated on 10% SDS-PAGE gels. After electrophoresis, separated protein bands were transferred into polyvinylidene fluoride membranes (Millipore, Cat. #IPVH00010) and blocked in 5% nonfat milk for 1 hour at room temperature. The membranes were then incubated with the primary antibodies at a recommend diluted ratio according to the manufacture instruction overnight at 4°C. After 3 times washing, membranes were incubated in 1:5000 horseradish peroxidase-linked secondary antibodies at room temperature for 2 hours. Finally, the membranes were washed for three times and were visualized using ECL Kit (Thermofisher, Cat. #34096).

For immunoprecipitation, cells were harvested and lysis in NP40 buffer, supplemented with PMSF and cocktail protease inhibitor. After centrifugation, supernatant was collected into fresh tubes. About 1mg of cell lysis was incubated with 1.0ug primary antibodies (anti-Flag, anti-DAB2IP, anti-p53, anti-HA) with rotation overnight at 4°C. Then 25ul protein A agarose beads (Sigma) were added into the lysates and a further incubation of 2 hours at 4°C with rotation was performed. Subsequently, breads were collected via centrifugation and washed 3-4 times with NP40 buffer. The immune-complex was released by boiling the breads in SDS loading dye and analyzed by western blotting.

### Quantitative real-time PCR (RT-qPCR)

Total RNA was extracted using TRIzol (Takara, Cat. #9108) and reverse-transcribed using PrimeScriptTM RT Master Mix reagent Kit (Takara, Cat. #RR036A) according to the manufacturer’s manual. Quantitative real-time PCR was carried out using TB Green™ Premix Ex Taq™ II Kit (Takara, Cat. #RR820A) in ABI 7300 Real-Time PCR System (Applied Biosystems, Foster City, CA). GAPDH gene expression was used as an endogenous control and the results from qRT-PCR were analyzed though the comparative Ct method (2-ΔΔCt). Primers’ sequences used in this research were provided as follow.

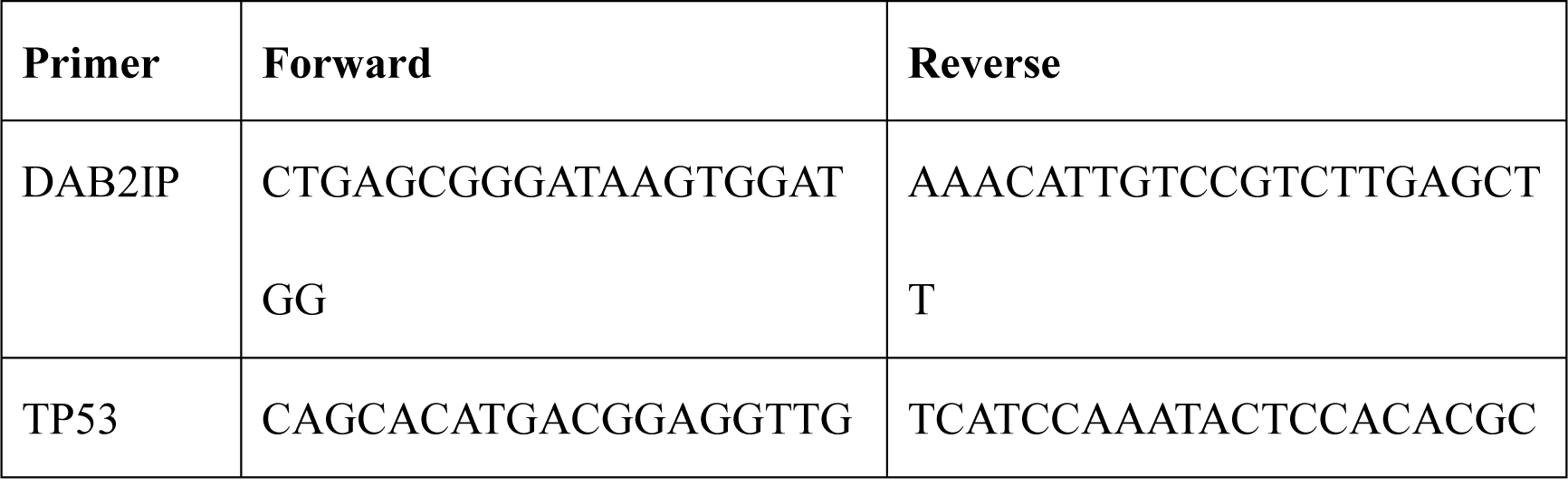
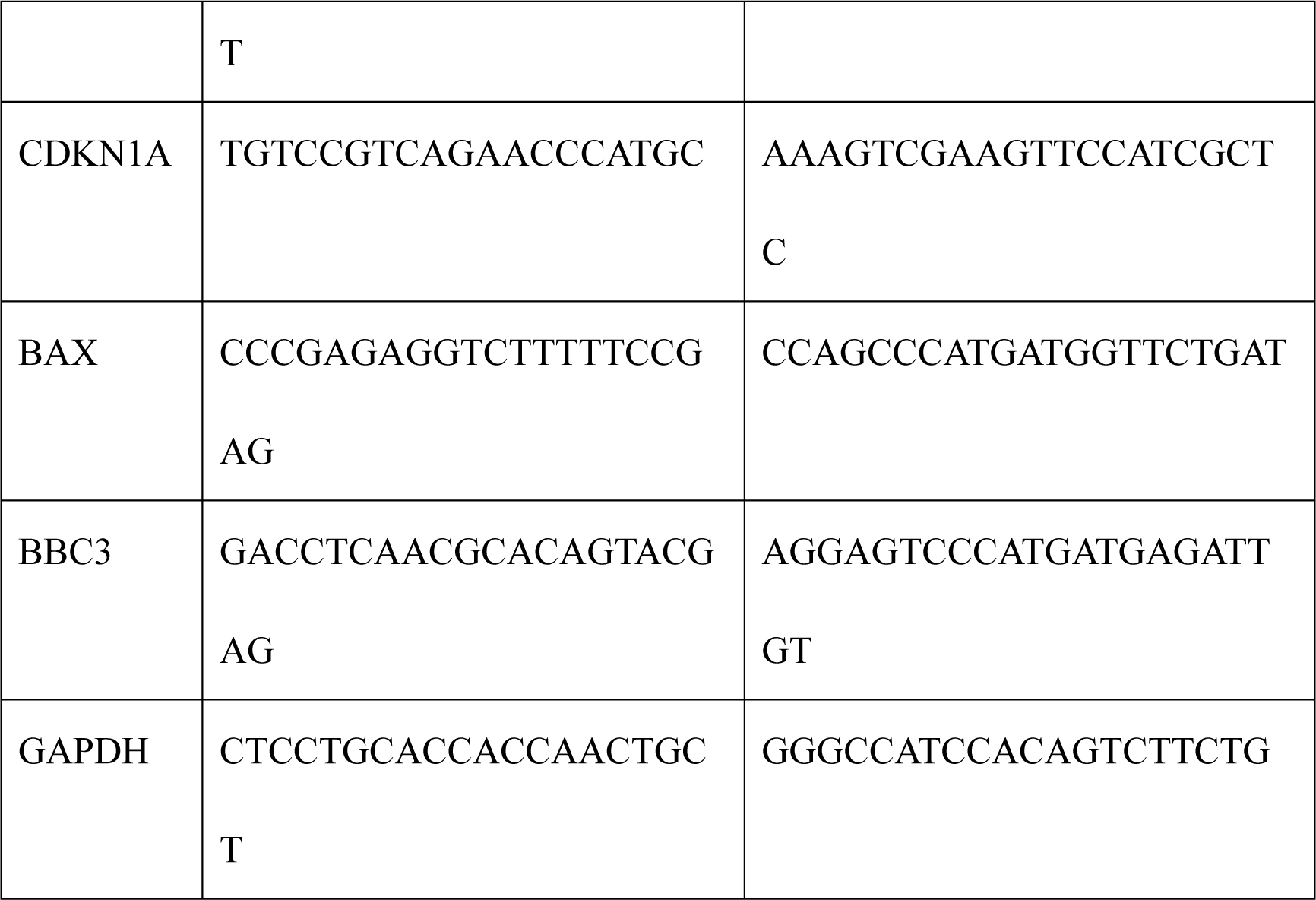

### Ubiquitination assay

DAB2IP or GRP75 was knocked down in SW48 cell lines via recombinant lentivirus or small interfering RNAs. HEK293T cells were stable transfected with pcDNA3.1-Myc-tagged-DAB2IP to overexpress DAB2IP expression. Then, these cell lines were co-transfected with pcDNA3.1-Flag-tagged-TP53 and HA-tagged-Ubiquitin plasmid. After 24-48 hours, cells were treated with MG-132 at a dose of 10uM for 8h and were lysed in NP40 buffer and centrifuged. The supernatants were collected and immunoprecipitated with anti-Flag agarose, and the immunocomplexes were examined using anti-HA antibody via western blotting.

### Mass spectrum analysis

The mass spectrum analysis was performed by Shanghai Lu-Ming Biotech Co., Ltd.

Briefly, protein was extracted via NP40 buffer added with cocktail protease inhibitor and separated on 10% SDS-PAGE gels. After electrophoresis, separated protein bands was strained with Coomassie brilliant blue. Next the protein bands were cut into 1mm^2^ gel particles and transferred into low protein binding tubes and rinsed twice with ultrapure water. Subsequently, gel was digested and the protein was purified according to the Katayama et al. (27).

For mass spectrum (MS) analysis, samples were pre-processed and loaded on the mass spectrometer (LTQ Orbitrap Velos Pro, Thermo Finnigan) according to the manufacturer’s protocol. Based on MS spectra, proteins were successfully identified based on 95% or higher confidence interval of their scores in the Proteome Discoverer 2.3 search engine (Thermo Scientific) and Uniprot-Homo database.

### Animals

All animal experiments were approved by Ethical Committee of Tongji Hospital. Briefly, female BABL/c athymic nude mice (age 4 w) were purchased from Charles River Inc., Beijing. The nude mice were subcutaneously injected 2 × 10^6^ tumor cells with Matrigel (Corning), 6 mice per group. After 4 weeks, the mice were euthanized, and the tumors were removed. After measuring the size, tumors were embedded, sliced and strained by hematoxylin-eosin (HE). The expression of DAB2IP and p53 was evaluated via immunohistochemistry. Ki-67 was used to evaluate the proliferation of tumor. TdT-mediated dUTP Nick-End Labeling assay kit (Biosci, Wuhan) was used to measured apoptosis rate of tumor cells.

### Public database and bioinformatics analysis

The gene expression dataset (GSE21510, (28)) of purified cancer cells in 104 patients with colorectal cancer was obtained from Gene Expression Omnibus database and was used in Gene Set Enrichment Analysis (GSEA) (http://software.broadinstitute.org/gsea/) (29). Mutation data, gene expression profile and prognostic data of colon adenocarcinoma patients was obtained from TCGA-COAD dataset (https://portal.gdc.cancer.gov). String database (http://www.string-db.org/) (30) was used for protein interaction analysis. Gene expression correlation was revealed by ’R2: Genomics Analysis and Visualization Platform (http://r2.amc.nl)’ using Sugihara dataset (28). Gene ontology enrichment were performed by DAVID software (https://david.ncifcrf.gov/tools.jsp). Mutation data of p53 in various colon cancer cell line was obtained from Cancer Cell Line Encyclopedia (CCLE, Broad Institute, https://portals.broadinstitute.org/ccle).

### Statistical analysis

Statistical analysis was performed by SPSS software package (version 19.0 for Windows; IBM, USA) and Prism 5 (GraphPad). In all graph, continuous data were expressed as mean ± standard deviation (SD), and statistically analyzed with student t test (two-tailed) or Analysis of Variance (ANOVA). A p value less than 0.05 was considered statistically significant.

## Results

### DAB2IP was positively associated with a better prognosis of colon cancer patients expressing wild-type TP53

To assess the potential correlation between the DAB2IP and TP53 protein levels in colon cancer, we evaluated a tissue chip containing 78 clinical specimens of malignant colon tumors via IHC staining. As shown in Figure 1A, the IHC score of TP53 was significantly increased in patients with high DAB2IP expression (Figure 1A). Additionally, gene set enrichment analysis (GSEA) of a public colorectal cancer expression profile (GSE21510) also revealed that p53 signaling was positively associated with DAB2IP expression (Figure 1B). R2 gene correlation analysis (http://r2.amc.nl) showed that DAB2IP was positively associated with TP53 target gene (CDKN1A, BAX, and PUMA) expression; however, no correlation was observed between DAB2IP and TP53 expression in colon cancer specimens (Figure 1C). Considering the frequency of TP53 mutations in colon cancer, we also analyzed the DAB2IP expression level in subgroups with wild-type TP53 and mutated TP53 from the TCGA-COAD dataset. As shown in Figure 1D, no difference in DAB2IP expression was observed between the two groups. These results indicated that DAB2IP could positively regulate wild type TP53 signaling at the posttranscriptional level.

**Figure 1.**
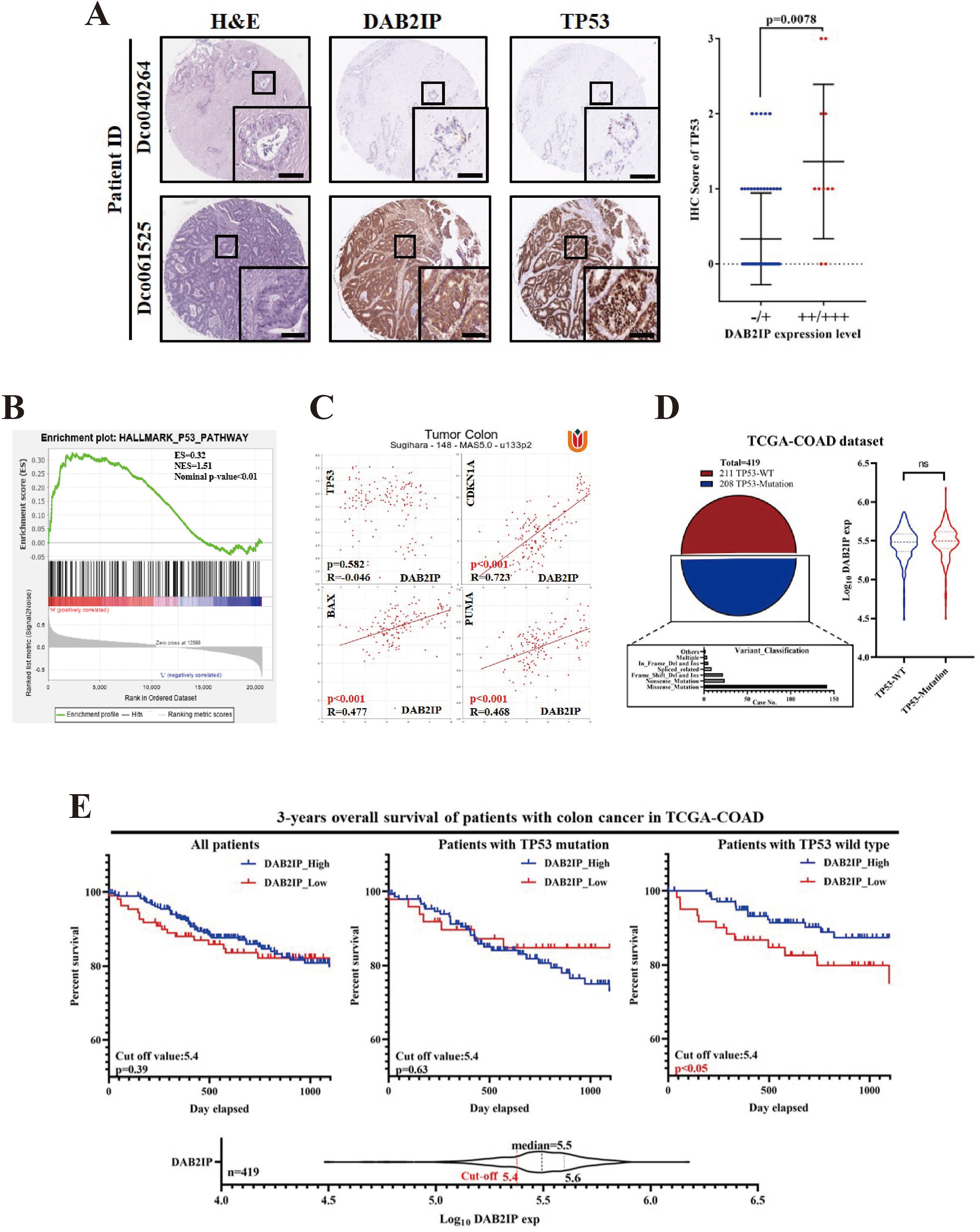
DAB2IP was positively associated with a better prognosis in colon cancer patients with wild-type TP53 expression. A) DAB2IP and TP53 expression in a tissue chip containing 78 clinical specimens of malignant colon tumor. IHC score of TP53 was significantly increased in tumor with DAB2IP high expression. (H&E: hematoxylin-eosin staining, p value was calculated via Students’ t test, Scale bar: 50 um). **B)** Gene set enrichment analysis (GSEA) of public colorectal cancer expression profile (GSE21510). Representative GSEA plots indicated that the p53 pathway was positively associated with DAB2IP high expression among 50 hallmark of cancer gene sets (NES=1.51, NOM p-value < 0.01). **C)** R2 gene correlation analysis of public clinical colorectal cancer dataset (Sugihara, n=148) revealed that DAB2IP was positively associated with TP53 target genes CDKN1A (R=0.723, p<0.001), BAX (R=0.477, p<0.001, PUMA(R=0.468, p<0.001), however, no correlation was observed between DAB2IP and TP53 expression (R=-0.046, p=0.582). **D)** TP53 mutation data and DAB2IP expression profile of colon cancer was obtained from TCGA-COAD dataset (n=419). Among them, 211 cases represented wild type TP53, while TP53 mutation was detected in the remaining 208 cases. Missense was the most common type in TP53 mutation. No difference of DAB2IP expression was observed between TP53 wild type and TP53 mutation subgroup (WT: wild type; Del: deletion; Ins: insertion; ns: no significance; p value was calculated via Student’s t test). **E)** Kaplan-Meier overall survival analysis based on DAB2IP mRNA expression in 419 colon cancer patients from TCGA-COAD dataset (211 with TP53 wild type and 208 with TP53 mutation). Cut-off value of DA2IP expression was lower quartile. Patients with high DAB2IP expression had significantly better prognosis than those with low DAB2IP expression in TP53 wild type subgroup (p<0.05).

To explore the effect of DAB2IP on colon cancer prognosis, we performed Kaplan-Meier analysis on the TCGA-COAD dataset. Interestingly, although high patients with high DAB2IP expression had a better prognosis than those with low DAB2IP expression in the wild-type TP53 subgroup (Figure 1E). These data suggested that DAB2IP could be positively associated with a better prognosis in colon cancer patients expressing wild-type TP53.

### DAB2IP exerted tumor-suppressive effects and positively regulated the TP53 protein levels in colon cancer cell lines expressing wild-type TP53

To evaluate the effects of DAB2IP on wild-type TP53 colon cancer, we first examined DAB2IP and p53 expression pattern in 8 colon cancer cell lines (CaCo2, HT29, HCT116, KM12, LoVo, SW48, SW480 and SW620) (Figure 2A). The western blot results showed that the DAB2IP protein was relatively absent in Lovo cells, while TP53 was relative low expression in Lovo and absent in HT29 cell (Figure 2A, up). Then, to assess the TP53 mutation type, the gene mutation data of the colorectal cancer cell lines were obtained from the CCLE database and analyzed (Figure 2A, down and Supplemental Figure 1). As shown in Figure 2A, HCT116 and SW48 cells expressed both DAB2IP and wild-type TP53. Thus, these two cell lines were selected for further studies.

**Figure 2.**
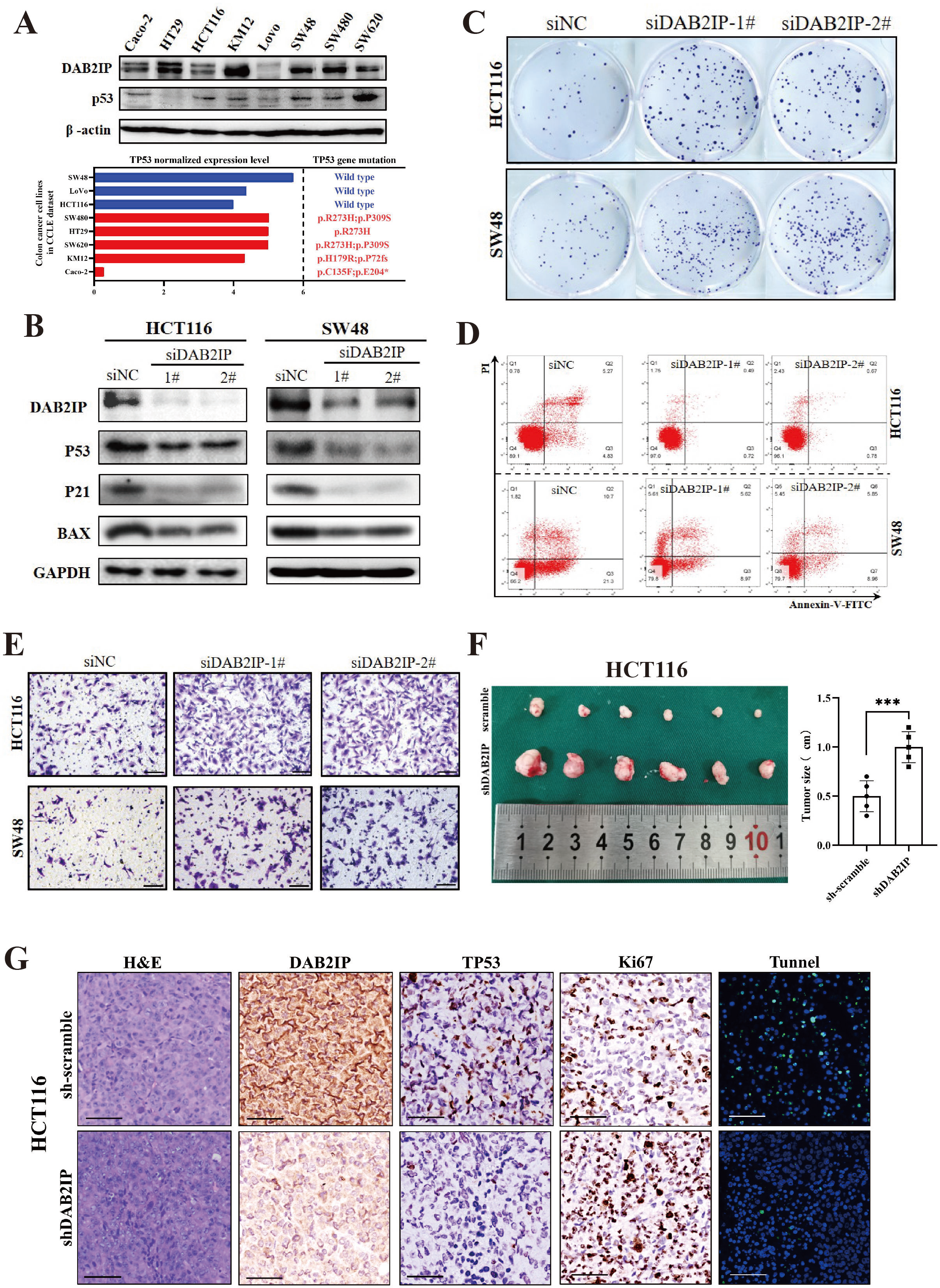
DAB2IP represented tumor-suppressive characteristics and positively regulated TP53 protein level in wild type TP53 colon cancer cell lines. **A)** Up: DAB2IP and p53 protein level was evaluated 8 colorectal cancer cell lines (Caco-2, HT29, HCT116, KM12, Lovo, SW48, SW480 and SW620) via western blot assay. Down: TP53 mutation data of the 8 colorectal cancer cell lines (Caco-2, HT29, HCT116, KM12, Lovo, SW48, SW480 and SW620) was obtained of CCLE database. TP53 in HCT116, Lovo and SW48 maintained wild type. **B)** DAB2IP was knocked down via siRNAs in both HCT116 and SW48 cell lines. Western blot assay showed that knockdown DAB2IP could significantly reduce the protein level of p53 and its downstream (p21 and BAX). **C)** Knockdown DAB2IP could significantly induce tumor cell clone formation in both HCT116 and SW48 cell lines. **D)** Knockdown DAB2IP could significantly suppress cell apoptosis in both HCT116 and SW48 cell lines. **E)** Knockdown DAB2IP could significantly induce tumor cell migration in both HCT116 and SW48 cell lines (scale bar: 100um). **F)** sh-scramble-HCT116 and sh-DAB2IP-HCT116 cells were subcutaneously injected into the Nude Mice. After 4 weeks, the mice were euthanized and the tumors were removed and measured. Representative images were shown. Tumors of mice injected with shDAB2IP-HCT116 were significantly larger than that of the sh-scramble mice. ***p<0.01 with Student’s t test. **G)** HCT116 xenografted with shDAB2IP exhibited higher levels of proliferation and lower levels of apoptosis than the control tumors. DAB2IP and p53 expression were detected via immunohistochemistry, cell proliferation and apoptosis were detected via Ki-67 straining and TUNEL assay. Representative images were shown. Scale bar: 50um.

According to previous results, the protein level of p53 was positively correlated with DAB2IP in clinical colon cancer specimens. To further elucidate whether DAB2IP could regulate p53 expression in vitro, we used siRNAs to knockdown DAB2IP in both the HCT116 and SW48 cell lines. Western blot assays revealed that knockdown of DAB2IP significantly reduced the protein level of p53 and its downstream targets (p21 and BAX) (Figure 2B). In contrast, overexpressing DAB2IP via the pcDNA3.1-DAB2IP plasmid in these two cell lines significantly increased TP53 expression (Supplemental Figure 2A). Additionally, knockdown of DAB2IP significantly induced tumor cell colony formation, inhibited cell apoptosis and promoted invasiveness and in both the SW48 and HCT116 cell lines (Figure 2C, D, E). Interesting, in colon cancer cell lines with mutated TP53 expression (SW480 and SW620, Figure 2A, down), neither knocking down nor overexpressing DAB2IP could affect the protein level of TP53 and its downstream (Supplemental Figure 2B, C). Thus, we speculated that DAB2IP could regulated wild-type TP53 expression.

To further investigated whether DAB2IP exerted oncogenic effects in vivo, we stably transfected HCT116 and SW48 cell line with sh-scramble or sh-DAB2IP lentivirus and subcutaneously transplanted it into nude mice. After 4 weeks, the mice were euthanized, and the tumor size was measured. Xenografted tumors from mice injected with shDAB2IP-HCT116 and shDAB2IP-SW48 were larger than those of sh-scramble mice (Figure 2F and Supplemental Figure 2D). Additionally, tumors with shDAB2IP exhibited higher levels of proliferation and lower levels of apoptosis than the control tumors (Figure 2G and Supplemental Figure 2E). These results indicated that the tumor-suppressive effects of DAB2IP in colon cancer cell lines expressing wild-type p53 and revealed a potential positive mechanism of DAB2IP regulation at the wild-type p53 protein level.

### DAB2IP inhibited the degradation of the p53 protein in a ubiquitin-proteasome-dependent manner

To elucidate the regulatory mechanism of DAB2IP-induced p53 expression, we first stably knocked down DAB2IP in the SW48 cell line with an shDAB2IP plasmid (DAB2IP-KD) (Figure 3A) and validated the mRNA level of TP53 and its downstream genes (CDKN1A, BAX and BBC3) through RT-qPCR. As shown in Figure 3B, knocking down DAB2IP significantly reduced the expression of the downstream genes of TP53 (CDKN1A, BAX and BBC2); however, DAB2IP did not exert a significant effect on the TP53 mRNA levels. Thus, we speculated that the regulation of DAB2IP-induced p53 expression might occur at the posttranscriptional stage.

**Figure 3.**
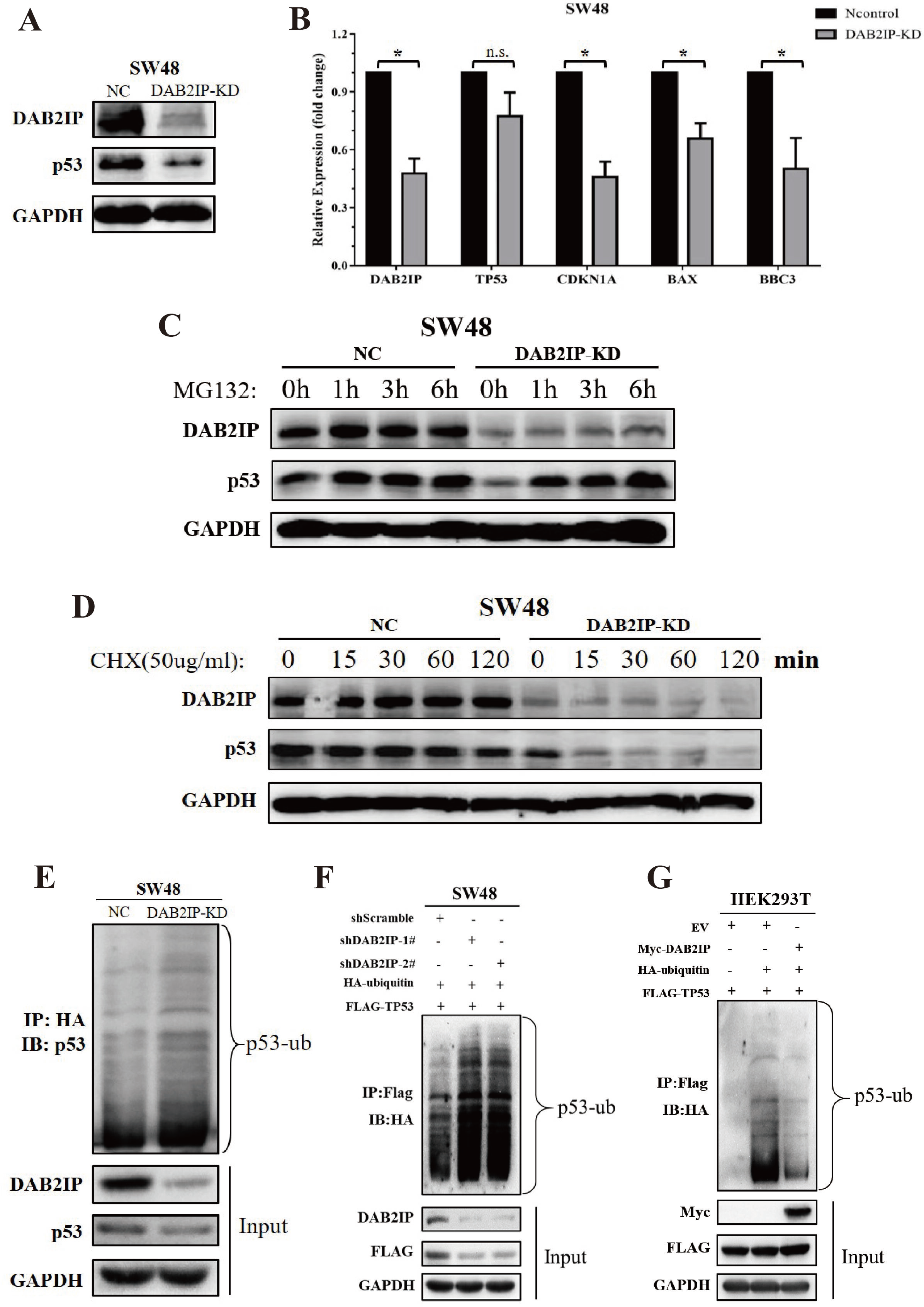
DAB2IP inhibited the degradation of p53 protein in a ubiquitin-proteasome-dependent manner. A) DAB2IP protein expression was stably knocked down in SW48 cells (labelled as DAB2IP-KD). Western blot assay showed that p53 protein level was significantly down-regulated in DAB2IP-KD group. **B)** RT-qPCR showed that knocking down DAB2IP could significantly reduce the expression of TP53 downstream genes (CDKN1A, BAX and BBC3), however, DAB2IP did not exhibit a significant influence on TP53 mRNA level. *p<0.05 by student’s t test; n.s. no significant difference. **C)** SW48 (control or DAB2IP-KD) was treated with MG132 at 10uM for the indicated time points. DAB2IP and p53 expressions were detected using western blot assay. **D)** SW48 (control or DAB2IP-KD) was treated with CHX at 150ug/ml for the indicated time points. DAB2IP and p53 expressions were detected using western blot assay. **E)** SW48 (control or DAB2IP-KD) was transfected with HA-ubiquitin plasmid and treated with MG132 at 10uM. HA antibody was used to immunoprecipitated ubiquitination proteins, and endogenous p53 in the immunocomplexes were detected via western blot assay. **F)** SW48 was co-transfected with HA-ubiquitin, Flag-TP53 plasmids, and infected with shDAB2IPs lentivirus, then treated with MG132 at 10uM for 8h. Flag antibody was used to immunoprecipitated exogenous p53 proteins, and ubiquitination p53 in the immunocomplexes were detected by HA antibody via western blot assay. **G)** HEK293T was co-transfected with HA-ubiquitin, Flag-TP53, and myc-DAB2IP plasmids, and treated with MG132 at 10uM for 8h. Flag antibody was used to immunoprecipitated exogenous p53 proteins, and ubiquitination p53 in the immunocomplexes were detected by Flag antibody via western blot assay.

Based on the previous results, we hypothesized that DAB2IP could regulate p53 at the protein synthesis/degradation level. To further explore the mechanism underlying DAB2IP-induced p53 signaling, we first used MG132, a reversible proteasome inhibitor, to indirectly evaluate protein synthesis efficiency in vitro. As shown in Figure 3C, after inhibiting protein degradation, knockdown of DAB2IP did not decrease the accumulation of the TP53 protein in the SW48 cell line. Next, we detected the half-life of p53 when DAB2IP was knocked down. After knockdown of DAB2IP, SW48 cells were treated with cycloheximide (CHX) at a dose of 50 µg/ml. The half-life of endogenous p53 was significantly shorter in the DAB2IP-KD SW48 cells than in the control cells (Figure 3D). These results indicated that DAB2IP acted as a negative regulatory factor on TP53 degradation.

The ubiquitin-proteasome pathway has been proven to be a highly effective pathway of p53 protein degradation in eukaryotic cells (31). To determine whether DAB2IP could regulate the ubiquitin-mediated degradation of the p53 protein, we first used a ubiquitination assay to detect the ubiquitination level of endogenous p53 protein in the SW48 cell line. As shown in Figure 3E, the ubiquitination level of endogenous p53 protein was significantly increased in DAB2IP-KD SW48 cells compared with control cells (Figure 3E). Subsequently, SW48 cells were coinfected with shDAB2IP lentivirus and Flag-tagged TP53 plasmid. The results showed that the ubiquitination of exogenous wild-type p53 protein was stronger in the DAB2IP-knockdown group than in the control group (Figure 3F). Similarly, the overexpression of DAB2IP also significantly inhibited exogenous p53 ubiquitination in the HEK293T cell line (Figure 3G). These results revealed that DAB2IP could inhibit the degradation of the p53 protein in a ubiquitin-proteasome-dependent manner.

### DAB2IP suppressed p53 ubiquitination through competitive binding GRP75

Previous results demonstrated that DAB2IP inhibited ubiquitin-proteasome-dependent p53 degradation. How DAB2IP regulates p53 ubiquitination is still unclear. In this section, we focused on exploring the detailed mechanism of DAB2IP-mediated TP53 ubiquitination.

First, a coimmunoprecipitation assay revealed that neither endogenous nor exogenous DAB2IP protein interacted with p53 directly (Supplemental Figure 3A, B). To screen for any potential mediator between DAB2IP and p53, Immunoprecipitation and mass spectrometry were performed and 166 proteins that interacted with both DAB2IP and p53 were identified in the SW48 cell line (Figure 4A, and Supplemental Figure 3C). Among them, TUBB, HSP90AA1, HSP90AB1, HSPA1B, HSPA5, HSPA8, XRCC5, GRP75 and RPL23 were classified into the “ubiquitin protein ligase binding” category via gene ontology enrichment (Figure 4B). Through the String protein interaction analysis (string-db.org), GRP75 (also known as HSPA9) and HSP90AA1 appeared to be a potential inhibitor of p53 protein (Figure 4C). Considering the relative higher binding capacity between DAB2IP and TP53 (Figure 4B), GRP75 were selected for further studies.

**Figure 4.**
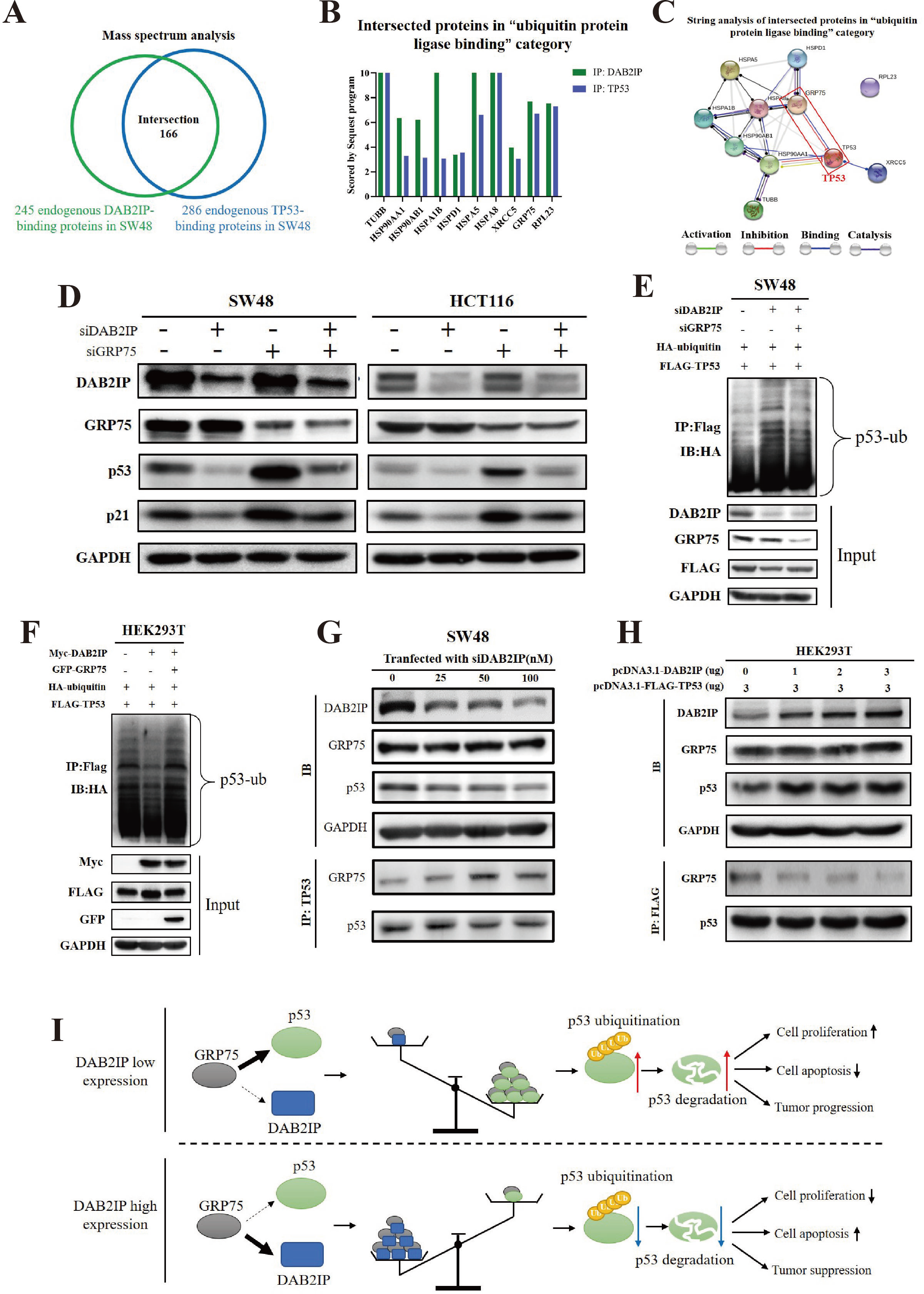
DAB2IP suppressed p53 ubiquitination through competitive binding GRP75. A) 245 endogenous DAB2IP-binding proteins and 286 endogenous p53-binding protein were detected via mass spectrum in SW48 cell line, among them, 166 proteins interacted with both DAB2IP and p53 protein. **B)** Proteins that interacted with both DAB2IP and p53 from (A) were used in gene ontology enrichment. Ten proteins involved in “ubiquitin protein ligase binding” were shown. Bar plot represented the Sequest score of these proteins. **C)** Using STRING analysis to identify the interaction between p53 and the ubiquitin protein ligase binding proteins from (B). GRP75 (also known as HSPA8) presented the potential inhibitor of p53. **D)** Knocking down GRP75 could significantly increase p53 and its downstream, p21, in SW48 and HCT116 cells with DAB2IP low expression. As determined by western blot assay. **E)** SW48 was co-transfected with HA-ubiquitin and Flag-TP53 plasmids, then knocked down DAB2IP and/or GRP75 with relevant siRNAs. After treated with MG132 at 10uM for 8h, Flag antibody was used to immunoprecipitated exogenous p53 proteins, and ubiquitination p53 in the immunocomplexes were detected by HA antibody via western blot assay. **F)** HEK293T was co-transfected with HA-ubiquitin and Flag-TP53 plasmids, then overexpressed DAB2IP and/or GRP75 with relevant plasmids. After treated with MG132 at 10uM for 8h, Flag antibody was used to immunoprecipitated exogenous p53 proteins, and ubiquitination p53 in the immunocomplexes were detected by HA antibody via western blot assay. **G)** After transfecting siDAB2IP into SW48 at the indicated concentrations, p53 antibody was used to immunoprecipitated endogenous p53 and its binding proteins. GRP75 in the immunocomplexes were detected by relevant antibody via western blot assay. **H)** After transfecting pcDNA3.1-DAB2IP and pcDNA3.1-FLAG-TP53 plasmids into HEK293T at the indicated concentrations, FLAG antibody was used to immunoprecipitated exogenous p53 and its binding proteins. GRP75 in the immunocomplexes were detected by relevant antibody via western blot assay. **I)** Model for how the DAB2IP regulates p53 expression and suppresses tumor progression.

To further elucidate the role of GRP75 in DAB2IP-regulated p53 degradation, we first used coimmunoprecipitation assays to verified that both endogenous DAB2IP and p53 could bind to GRP75 in the SW48 cell line (Supplemental Figure 3D, E). Western blotting assay revealed that GRP75 could negatively regulate p53 and its downstream at the protein level in both SW48 and HCT116 cell lines (Supplemental Figure 3F). Subsequently, we found that knocking down GRP75 could increase the p53 protein levels in cells with low DAB2IP expression and significantly reversed DAB2IP-regulated clone formation and cell apoptosis in both the SW48 and HCT116 (Figure 4D and Supplemental Figure 3G, H). Through a ubiquitination assay, we observed that knockdown of GRP75 could significantly reverse DAB2IP-mediated p53 ubiquitination in the SW48 cell line (Figure 4E). Similarly, overexpressing GRP75 also increased the ubiquitination level of exogenous wild-type p53 in the HEK293T cells transfected with the DAB2IP plasmid (Figure 4F). These findings indicated that GRP75 acted as a positive regulator in DAB2IP-mediated p53 ubiquitination.

Previous results revealed that both DAB2IP and p53 could bind to GRP75 (Supplemental Figure 3D, E). Considering the promoting effect of GRP75 on DAB2IP-mediated p53 ubiquitination, we hypothesized that DAB2IP could suppress ubiquitination-mediated p53 degradation by competitively binding to the GRP75 protein. To verify this hypothesis, we transfected siDAB2IP into SW48 cells at the indicated concentrations. Immunoprecipitation revealed that with a decline in DAB2IP protein levels, the binding capacity of GRP75 to p53 was gradually increased (Figure 4G). Similarly, the amount of GRP75 bound to p53 was negatively correlated with the DAB2IP protein levels in HEK293T cells (Figure 4H). These data revealed that DAB2IP suppressed p53 ubiquitination by competitive binding to the GRP75 protein.

In summary, by competitive binding to GRP75, DAB2IP inhibits GRP75-mediated p53 ubiquitin-degradation, thus increasing the p53 protein levels, promoting p53 signaling, and finally exhibiting tumor-suppressive characteristics against colon cancer (Figure 4I).

## Discussion

The important roles of p53 in tumor suppression, including cell death, damage repair, metabolism, proliferation and senescence, have been widely established in cancer development (8, 12). Due to its frequent alterations in human cancer, the mutation state of the TP53 gene has become a highly valuable predictive and prognostic biomarker in multiple tumors (4, 32). The mutation incidence of TP53 is nearly 40% in colon cancer, while the remaining 60% of patients express the wild-type protein (33). Numerous studies have focused on exploring the oncological role of mutant p53; however, the mechanism of genesis and progression of tumors expressing wild-type p53 is still unclear. In the current study, through a combination of bioinformatics analysis and clinical specimens, we unexpectedly revealed that DAB2IP had a higher anti-oncologic effect in colon cancer patients expressing wild-type p53, and it positively regulated p53 signaling by influencing its protein level.

DAB2IP was first identified as a DOC-2/DAB2 interactive protein and was demonstrated to be a tumor suppressor of cell growth, metastasis and other aspects of cancer progression by our group (34). As a scaffold protein, DAB2IP contains multiple domain structures to interact with key components of various pathways, and then, it regulates tumor-related signaling, such as RAS/ERK (35), ASK1/JNK (20), PI3k/Akt (19), androgen receptor (36), NF-kb (24), and b-catenin/TCF signaling (22). Previous studies have demonstrated that mutant p53 can bind to DAB2IP and interfere with its function in Akt activation and TNF signaling (25, 26). Additionally, Lunardi A. et al. also reported that DAB2IP can bind to wild-type p53, at least in vitro and in transient overexpression conditions, although whether such an interaction could occur physiologically is still unclear (37). Our study is the first to discover the positive regulatory effect of DAB2IP on wild-type p53 and its downstream protein levels in colon cancer cells. Contrary to earlier reports, DAB2IP may act upstream of the wild-type p53 protein and inhibit tumor progression by directly activating p53 signaling. Our findings present a new possibility regarding the mechanism of the DAB2IP-induced tumor suppression effect in colon cancer.

Subsequently, our research elucidated how DAB2IP regulated wild-type p53 expression. One of the key mechanisms by which p53 is regulated is through rapid degradation via the ubiquitin proteasomal pathway (31). Coincidentally, our present study also revealed that DAB2IP could regulate p53 abundance at the posttranscriptional levels. Mechanistically, DAB2IP significantly inhibited p53 ubiquitination, prolonged the half-life of wild-type p53, and activated p53 signaling. These findings revealed the function of DAB2IP in p53 ubiquitin-proteasome-dependent degradation and increased our understanding of DAB2IP-mediated tumor suppression.

Through an in vitro assay, we found that DAB2IP could not directly interact with the p53 protein. Additionally, DAB2IP did not exhibit E3 ligase activity, which is necessary for the selection of proteins for ubiquitination-dependent degradation (34, 38). Thus, we suggested that an intermediate should exist between the two proteins and be involved in the regulation of DAB2IP-mediated p53 ubiquitination.

To date, nearly 20 E3 ligases (including MDM2, CHIP, TRIMs, etc.) and 8 non-E3 ligases that target p53 for proteasomal degradation have been identified (31). Among them, MDM2 has been regarded as the canonical regulator of p53 degradation, while CHIP, an E3 ligase able to target its substrates for proteasomal degradation by binding to the C termini of Hsc70 and Hsp90 and mediating the ubiquitination of chaperone-bound proteins, has been widely reported to drive p53 ubiquitination in the noncanonical pathway (39). In mass spectrometry analysis, MDM2 was not detected among the DAB2IP binding proteins. Interestingly, GRP75, a chaperone protein that interacts with the CHIP E3 ligase (40), has been verified to bind to both endogenous and exogenous DAB2IP proteins and to play an important regulatory role in DAB2IP-mediated p53 ubiquitination. Thus, we assumed that DAB2IP regulated p53 protein levels via ChIP-dependent ubiquitination.

GRP75, also known as mortalin and HSPA9, is a member of the family of glucose-regulated proteins, which belong to the stress-inducible molecular chaperones of the heat shock protein (HSP) family and play an important role in the regulation of protein quality control and degradation (41). Early studies showed that GRP75 could induce abrogated cytoplasmic retention of wild-type p53 (42) and facilitated proteasome-mediated degradation of p53 by the ChIP-GRP75 complex (43). Additionally, disrupting the formation of the GRP75-CHIP complex could consequently suppress the CHIP-mediated destabilization of p53 (43). In our study, we first revealed that DAB2IP could suppress p53 ubiquitination by competitively binding to the GRP75 protein. These findings revealed a new component of CHIP-mediated p53 degradation and might increase our understanding of the dynamic regulation of p53 stabilization.

In summary, our study presents a novel function of DAB2IP in GRP75-driven wild-type p53 ubiquitination-degradation. This finding not only reveals a new signaling axis that is involved in DAB2IP-induced tumor suppression but also provides a novel molecular aspect of the p53 pathway.

## Acknowledgements

We thank Jer-Tsong Hsieh for providing DAB2IP overexpressing plasmids. We thank experimental medicine center of Tongji Hospital for providing support to our experiment, and Department of Pathology of Tongji Hospital for immunohistochemistry analysis. We apologize to the colleagues whose work was not cited due to space constraint.

## Supplementary Information

**Supplemental Figure 1.**
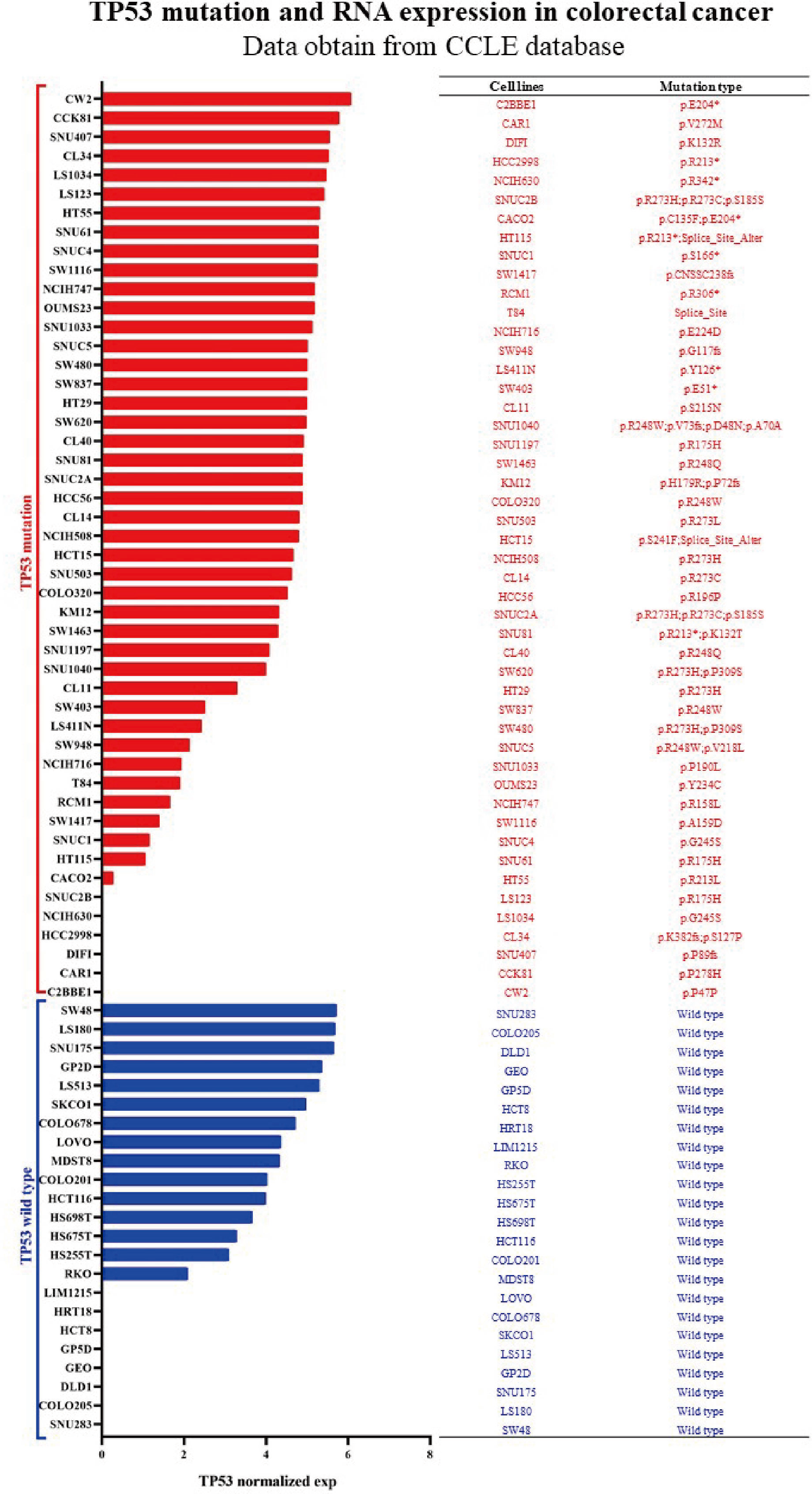
TP53 mRNA expression level and mutation type in colorectal cancer cell lines from CCLE database.

**Supplemental Figure 2.**
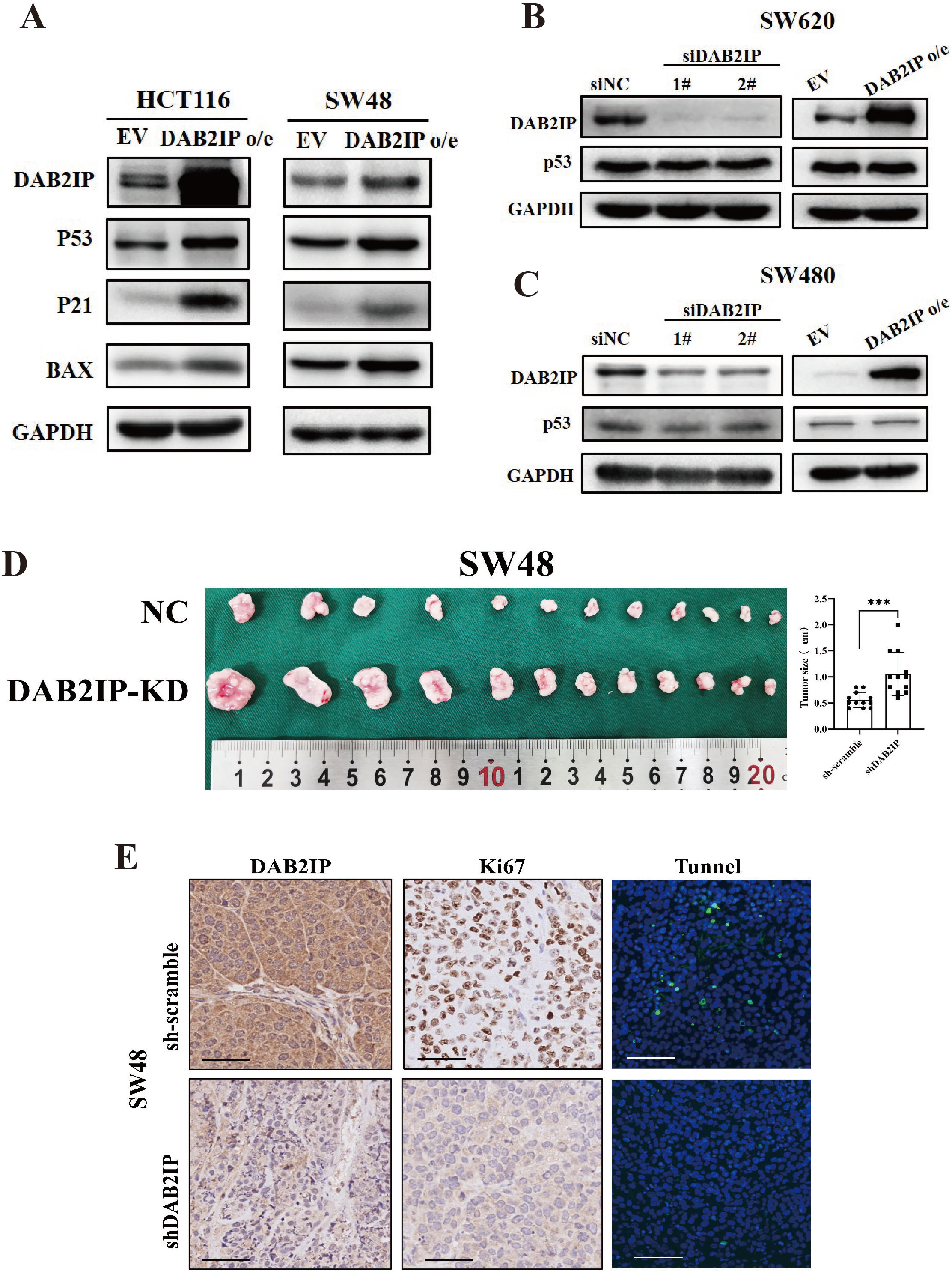
DAB2IP represented tumor-suppressive characteristics and positively regulated TP53 protein level in wild type TP53 colon cancer cell lines. **A)** Western blot showed that overexpression of DAB2IP could significantly increase the protein level of p53 and its downstream (p21 and BAX) in both SW48 and HCT116 cell lines. **B)** Western blot showed that neither knocking down nor overexpressing DAB2IP could affect the protein level of TP53 and its downstream in SW620 cell line. **C)** Western blot showed that neither knocking down nor overexpressing DAB2IP could affect the protein level of TP53 and its downstream in SW480 cell line. **D)** sh-scramble-SW48 and sh-DAB2IP-SW48 cells were subcutaneously injected into the Nude Mice. After 4 weeks, the mice were euthanized and the tumors were removed and measured. Representative images were shown. Tumors of mice injected with shDAB2IP-SW48 were significantly larger than that of the sh-scramble mice. ***p<0.01 with Student’s t test. **E)** SW48 xenografted with shDAB2IP exhibited higher levels of proliferation and lower levels of apoptosis than the control tumors. Representative images were shown. Scale bar: 50um.

**Supplemental Figure 3.**
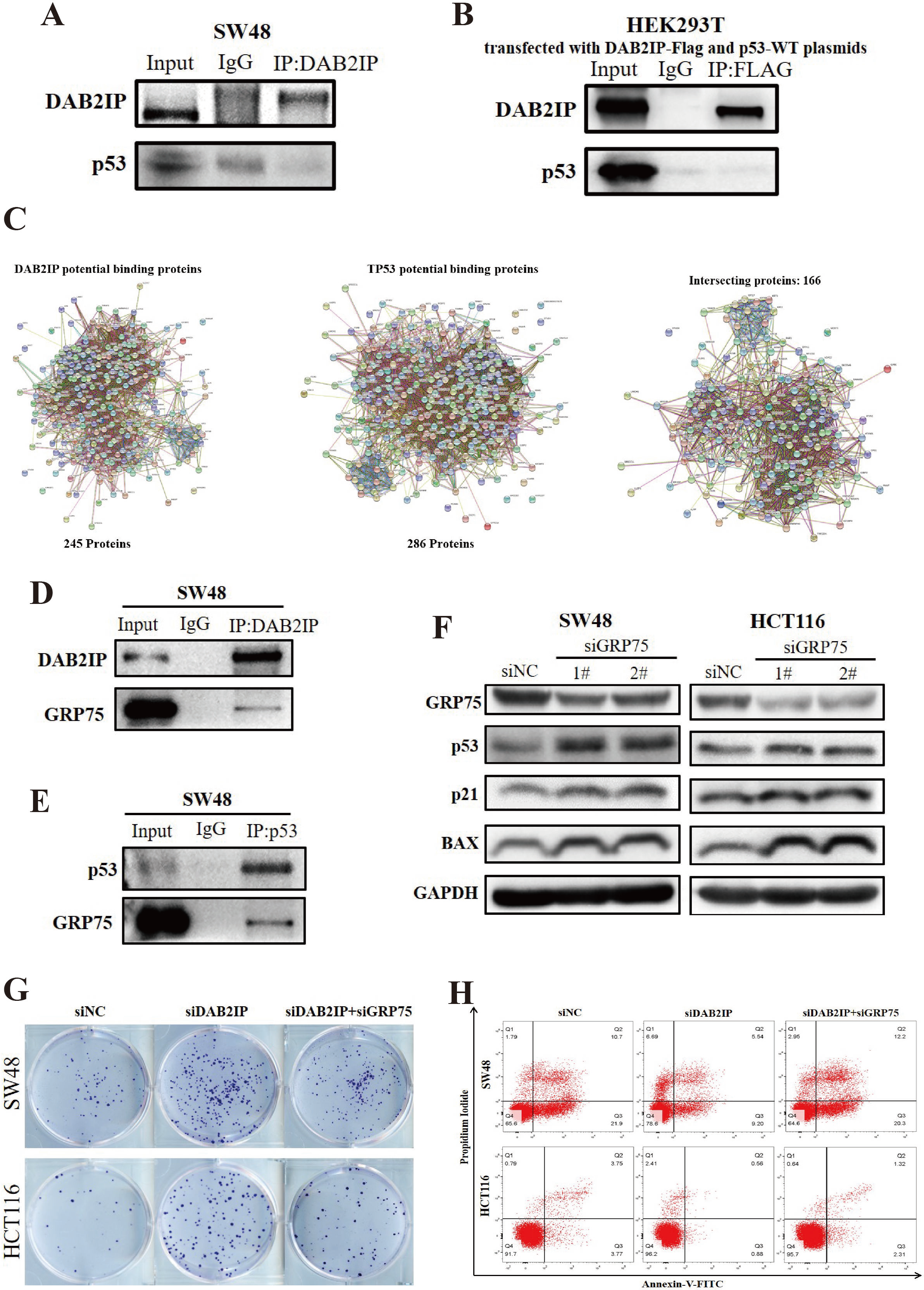
Screening the potential proteins that interacted with DAB2IP and TP53 protein. **A)** Endogenous DAB2IP could not immuno-precipitate p53 in SW48 cell line. **B)** Exogenous p53 could not bind with DAB2IP in HEK293T cell line. **C)** 245 endogenous DAB2IP-binding proteins and 286 endogenous p53-binding protein were detected via mass spectrum in SW48 cell line. Through intersecting the binding protein, 166 proteins that interacted with both DAB2IP and p53 protein were detected in SW48 cell lines. **D)** Coimmunoprecipitation assay revealed that endogenous DAB2IP could bind to GRP75 in SW48 cell lines. **E)** Coimmunoprecipitation assay revealed that endogenous p53 could bind to GRP75 in SW48 cell lines. **F)** Western blot assay showed that knockdown GRP75 could significantly increase the protein level of p53 and its downstream (p21 and BAX) in both SW48 and HCT116 cell lines. **G)** Knocking down GRP75 could significantly reduce clonal formation in both SW48 and HCT116 cells with DAB2IP low expression. **H)** Knocking down GRP75 could significantly induce cell apoptosis in both SW48 and HCT116 cells with DAB2IP low expression.

